# Tissue-Driven Hypothesis of Transcriptome Evolution: An Update

**DOI:** 10.1101/054890

**Authors:** Howard T. Hallmark, Jeffrey A Haltom, Xun Gu

## Abstract

In past decade, many reports have demonstrated that tissues in multi-cellular organisms may play important roles to shape the pattern of genome evolution. The tissue-driven hypothesis was then coined, claiming that tissue-specific factor as the common resource of functional constrain may underlie the positive correlations between tissue expression divergence, sequence divergence, or the expression tolerance of duplication divergence. However, the original version of tissue-driven hypothesis cannot rule out the tissue-specific effect of mutational variance. In this perspective, we solve this problem by modifying the evolutionary model that underlies the tissue expression evolution. Reanalysis of the microarray data reanalysis has revealed the relative importance between tissue-specific functional constraints and mutational variances in the tissue evolution. Finally, we outline how to utilize RNA-seq technology to further investigate the tissue expression evolution in the case of multiple tissues and species.

## Tissue expression evolution

In multi-cellular organisms, understanding the roles of tissue-specific factors during the course of genome evolution is the first step to investigate the emergence of biological complexity (Arendt 2008) but the tissue evolution remains obscure and controversial (Chan et al. 2009; Yanai and Hunter 2009). Thanks to the invent of high throughput technologies especially the microarray chips, a number of reports in the past decade have revealed interesting evolutionary patterns of tissue expression divergence in primates (Enard et al 2002; Gu and Gu 2003; Khaitovich et al. 2004a, 2004b; 2005a, 2005b, 2005c), in mammals (Duret and Mouchiroud 2000; Su et al. 2004; Gu and Su 2007), aswell as in fruitflies (Rifkin et al. 2003). The earliest work included Duret and Mouchiroud (2000) who showed that the rate of protein sequence divergence was negatively correlated with the tissue broadness of gene expression. In other words, broadly expressed proteins tend to evolve slowly, and *vice versa*. With the help of human Affymetric microarray chips, Enard et al. (2002) conducted a genome-wide expression analysis in the brains among primates, claiming a human lineage-specific acceleration of expression divergence. Follow-up analyses from our group (Gu and Gu 2003) and (Caceres et al. 2003; Uddin et al. 2004) suggested that up-regulation might be the major pattern during the evolution of human brain. Moreover, a detailed analysis on expression profiles in primates (Gilad et al. 2006) revealed a rapid evolution of human transcription factors.Together, these studies have provided an updated version of the regulatory hypothesis of human-chimpanzee split (King and Wilson 1975). Noticeably, Babbin et al. (2010) showed that both noncoding and protein-coding RNAs contributed to gene expression evolution in the primate brain, and Chodroff et al. (2010) showed the role of long noncoding RNA genes expressed in brains among diverse amniotes. Meanwhile, Khaitovich et al. (2005) addressed another important issue, that is, whether the rate of expression divergence among species differs among tissues. They analyzed genome-wide expression profiles in several tissues in primates, and observed that the brain or cerebellum tissue may have under stronger expression conservation than other tissues under study (testis, heart, liver and kidney); in particular, testis showed a rapid expression divergence.

## Stabilizing selection model for tissue-driven hypothesis

Gu and Su (2007) coined *the tissue-driven hypothesis*, postulating that tissue-specific factors may serve as the common functional constraints that may be imposed on different aspects of genome evolution. Under this framework, three specific predictions were derived: (*i*) a positive correlation between tissue expression distance and protein sequence distance between species; (*ii*) a positive correlation of tissue expression distance between species with that between duplicate genes; and (*iii*) tissue-specific factors and tissue broadness are two independent, additive recourses that shape the pattern of genome evolution. Initial analyses (Gu and Su 2007; Su and Gu 2007) have provided strong evidence for supporting these predictions. Indeed, most recent related studies can be explained by the tissue-driven hypothesis, in spite of various terminologies used by different authors (Brawand et al. 2011).

Yet we have recently realized a theoretical drawback of the tissue-driven hypothesis recently. Shortly speaking, tissue expression evolution is driven by two factors: the mutational variance accessible to the tissue, and the functional constraints of the tissue. In other words, variation of each factor among tissues would results in the variation of expression divergence among tissues, and the original version of tissue-driven hypothesis (Gu and Su 2007) did not distinguish between these two possibilities.

Using tissue expression distances (*E_ti_*) between human and mouse for 29 orthologous tissues, Gu and Su (2007) found a considerable variation of human-mouse expression divergence among tissues. As discussed above, there are two alternative mechanisms that can explain this observation: The first one invokes the effect of tissue-specific functional constraint (functionally important tissues tend to have strong constraints on expression divergence), and the second one invokes the effect of tissue-specific mutational variance (tissue-specific regulatory networks shape the consequence of regulatory mutations.

It has been argued that gene expression may be optimized by natural selection (Bedford and Hartl 2009). Gu and Su (2007) invoked the stabilizing selection model (Hansen 1997) to describe the tissue-specific selection constraint on the expression divergence; though many other models were proposed (Gu 2004; Eng et al. 2009). For a gene expressed in a certain tissue (*ti*), the stabilizing selection on the expression level *x* follows a Gaussian fitness function *f_ti_*(*x*) =exp[‐ *w_ti_*(*x*‐ θ)^2^], where θ is the optimal expression level, *w_ti_* is the coefficient of stabilizing selection on gene expression in tissue *ti*; a large *w_ti_* means a strong selection pressure, and *vice versa*. Under this model, the evolution of tissue expression follows an Ornstein-Uhlenback (OU) process, based on which the tissue expression distance is given by 
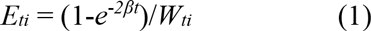

Where *W_ti_*= 2*N_e_W_ti_* is the strength of stabilizing selection against the expression divergence, and *N_e_* is the effective population size; β= *W_ti_* ε^2^ is decay-rate of expression divergence, and ε^2^ is the mutational variance.

Owe attempt to solve this problem, arguing that the revised tissue-driven hypothesis should include two sub-hypotheses: the tissue-specific functional constraint, as well as its alternative tissue-specific mutational variance (She et al. 2009). Moreover, extending the underlying evolutionary model allows further data exploration. Microarray data reanalysis has revealed their relative importance in shaping the pattern of tissue evolution. Finally, we discuss how to utilize novel next-generation technologies (NGS) such as RNA-seq to investigate tissue expression evolution when genome-wide expression data are available in multiple tissues and multiple species.

## Selection-mutation balance of tissue expression evolution

We solve this problem by formulating a stochastic model called the stationary Ornstein-Uhlenback (sOU) process model (Hansen 1997;Butler and King 2004). As shown in Material and Methods, the sOU model depends on two parameters: (i) The strength of tissue-specific functional constraint is measured by *W_ti_*; a large value indicates a strong constraint, and *vice versa*. It can be shown that tissue expression distance between twospecies is saturated to 1/*W_ti_* as the evolutionary time (*t*) is sufficiently large. And (ii) the mutational variance (2ε^2^*t*) measures the mutational capacity that drives the tissue-specific expression divergence between species. Moreover, the stationary assumption implies that the expression variance remains a constant during the expression divergence, under which we are able to estimate the two parameters.

In Eq.(1), we have only one observation (*E_ti_*) but two unknown parameters. To solve this problem, we need an additional equation that can be derived under the assumption of stationary OU process, which means that the expression variance remains invariant during the course of expression evolution. Let *R_ti_* be the coefficient of expression correlation between two species of the same tissue. Under the stationary assumption, it is given by 
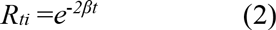

Hence, one can easily estimate *W_ti_* from two observations *E_ti_* and *R_ti_*, that is, by replacing *e*^−2β*t*^ in Eq.(B-l) with Rti according to Eq. (2), we have

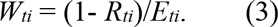

## Re-analysis of tissue-driven hypothesis

### Estimation of tissue-driven factors (Wti)

We first showed that the expression variance remained roughly the same between the human and mouse in each of 29 tissues, indicating that the assumption of selection-mutation balance holds approximately. For each tissue of two species (human and mouse), we estimated *W_ti_* from the observed tissue expression distance (*E_ti_*) and the coefficient of expression correlation (*R_ti_*) (Table 1). The first question one may ask is to what extent the variation of expression divergence among tissues can be explained by the tissue factor. Simple calculation shows that the tissue expression distance is negatively correlated with the tissue-specific functional constraint (*W_ti_*) (*R*^2^=0.57,*p*-value < 0.001). We therefore conclude that tissue-specific function constraint and tissue-specific mutational variance may explain equally the variation of expression distance among tissues.

**Table 1.** A summary of our estimates for the strength of functional constraints (*W*) and the distance of mutational variance (2ε2*t*)

| | $W$ | $2\varepsilon^2t$ |
| --- | --- | --- |
| Nerve system (7) | 0.640±0.026 | 2.619±0.580 |
| Reproduction system (4) | 0.521±0.091 | 3.105±0.553 |
| Soft tissues (8) | 0.495±0.033 | 2.456±0.151 |
| Endocrino-system (5) | 0.558±0.057 | 3.770±0.984 |
| Immuno-system (5) | 0.493±0.070 | 2.085±0.218 |
| all tissues (excluding nerves) (22) | 0.521±0.027 | 2.789±0.271 |
| all tissues (29) | 0.550±0.024 | 2.748±0.237 |

Note: According to the physiological roles, we tentatively classified 29 tissues into five groups: (i) nerve system, including cerebellum (*cb*), olfactory bulb (*oc*), amygdale (*ad*), hypothalamus (*hp*), trigeminal (*tm*), dorsal root ganglion (*dr*), and pituitary (*pi*); (ii) reproduction system, including testis (*ts*), uterus (*ur*), ovary (*ov*), and placenta (*pl*); (iii) soft-tissues, including heart (*ht*), kidney (*kn*), liver (*li*), lung (*lu*), adipose tissue (*at*), skeletal muscle (*sm*), tongue (*to*), and trachea (*tc*); (iv) endocrino-system, including adrenal gland (*ag*), pancreas (*pc*), prostate (*pt*), salivary gland (*sg*), thymus (*tm*), and thyroid (*tr*); and (v) immuno-system, including bone marrow (*bm*), CD4<sup>+</sup> T-cells (*T4*), CD8<sup>+</sup> T-cells (*T8*), and lymph node (*ln*).

In Table 1, 29 tissues are classified into several groups. We observed that neuro-related tissues, on average, have stronger tissue-specific function constraints (*W_ti_*) than neuro-unrelated tissues (*p*-value< 0.01, *t*-test), whereas no statistically significant difference in tissue-specific mutational variance (2ε^2^*t*) was found (*p*-value>0.10, *t*-test). Though we confirmed that several tissues have relaxed functional constraints, i.e., a low*W_ti_*, including testis (*W_ti_*=0.33), CD4 (*W_ti_*=0.36), CD8 (*W_ti_*=0.34) and pancreas (*W_ti_*=0.39), we found no significant difference in either *W_ti_* or 2ε^2^*t* among other biological systems except for the neuro-system.

### Decomposition of *E_ti_-D_ti_* (expression-sequence) correlation into *W_ti_-D_ti_* and 2ε^2^*t*-*D_ti_* correlations

Let *D_ti_* be the mean evolutionary distance of protein sequence (between the human and mouse) over a set of genes expressed in tissue *ti*. One important prediction of the tissue-driven hypothesis (Gu and Su 2007) is the existence of positive correlation between *E_ti_* and *D_ti_*, which reflects the common micro-environment of tissue (*ti*) on the expression divergence and protein sequence divergence, respectively. In Materials and Methods, we decomposed this prediction into the *W_ti_-D_ti_* correlation for the common tissue-specific functional constraints, and the 2ε^2^*t*-*D_ti_* correlation for tissue-specific mutational variance, respectively. Gu and Su (2007) considered two expression status of a gene in tissue *ti*, i.e., high expression or normal expression. In both cases, we indeed found a significant negative correlation between *W_ti_* and *D_ti_*. In the case of normal expression (Fig. 2, panel A), the coefficient of correlation is *r*=‐0.42 (*p*< 0.01), while *r*=‐0.31 (*p*< 0.01) in the case of high expression. Interestingly, we found that tissue-specific mutational variance showed a positive correlation with tissue protein distance (*r*=‐0.44, (*p*< 0.01 fornormally-expressed genes shown in Fig.2 panel B, and *r*= 0.38 for highly expressed gene, *p*< 0.01). We explain this positive 2ε^2^*t*-*D_ti_* correlation as the common tissue-specific genetic or epigenetic buffering against mutational effects.

**Figure 1.**
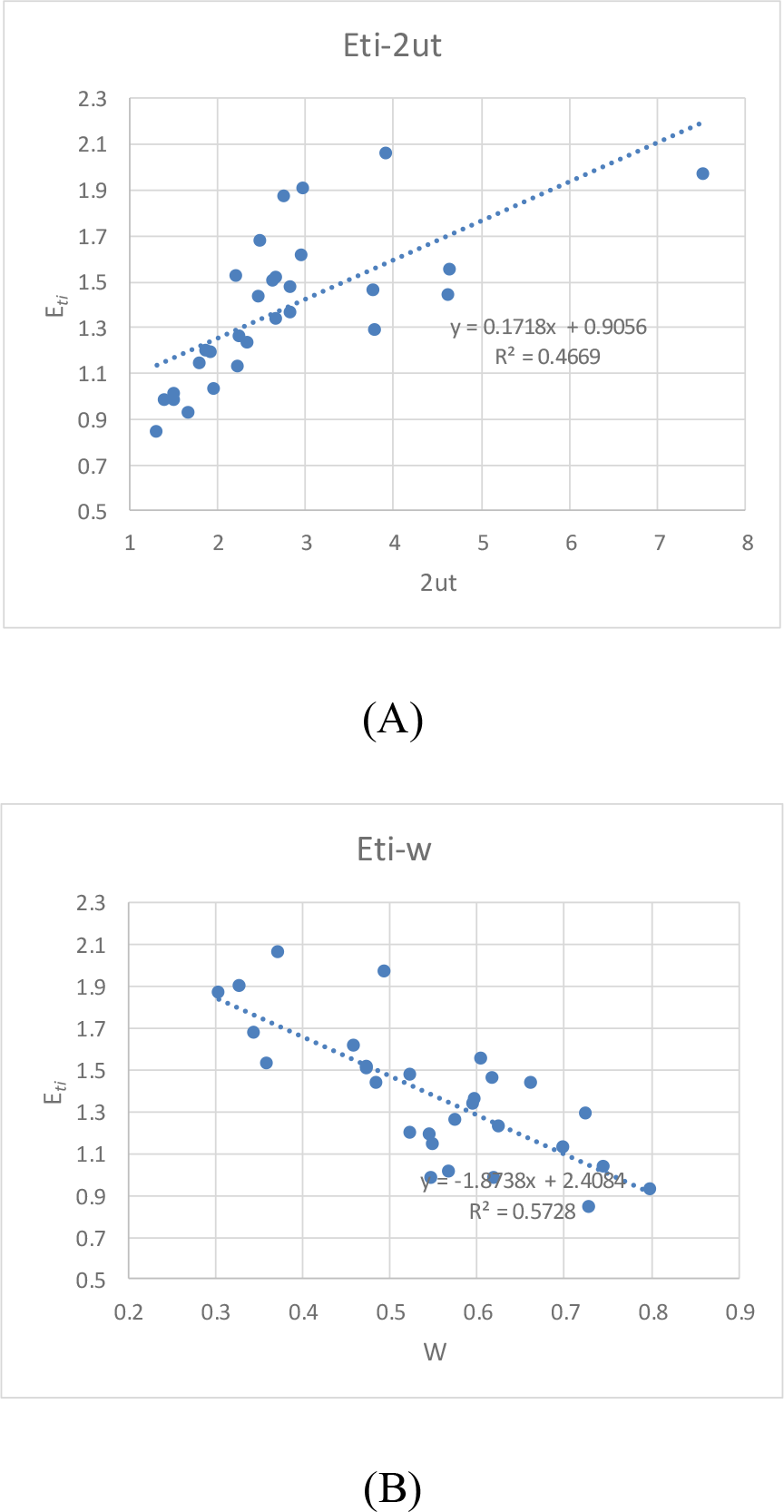
(A) Negative correlation between tissue expression distances (*E_ti_*) and tissue-specific function constraints (*W_ti_*) (*R*^2^=0.57, *p*-value<0.001). (B) Positive correlation between tissue expression distances (*E_ti_*) and tissue-specific mutation accessibility (2ε^2^*t*) (*R*^2^=0.47, *p*-value<0.001). (C) No statistically significant correlation between *W_ti_* and 2ε^2^*t* (*p*-value>0.25). See the footnote of Table 1 for the list of tissue names and abbreviations.

**Figure 2.**
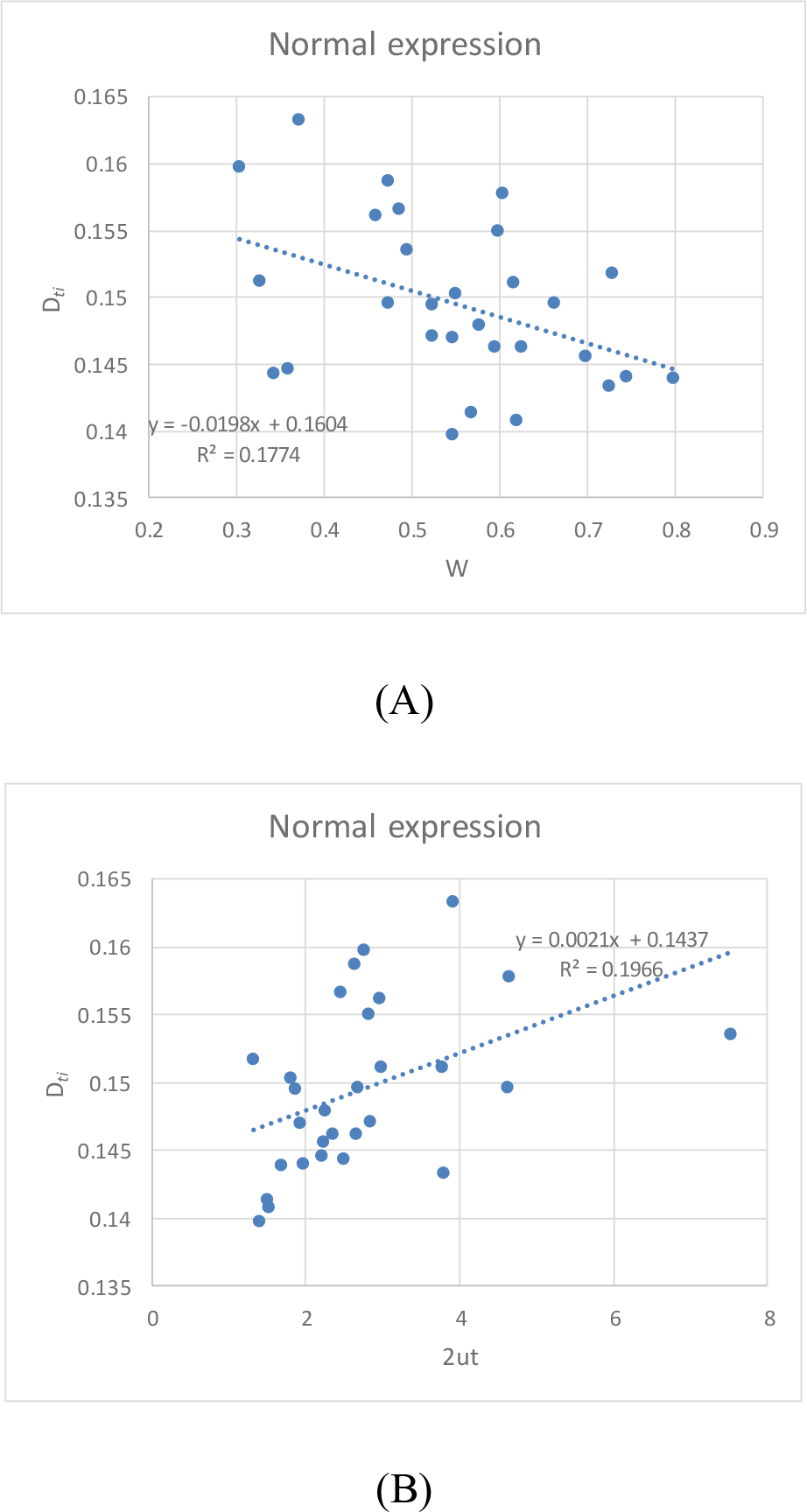
Negative correlation between tissue-specific function constraints (*W_ti_*) and tissue protein distance (*D_ti_*) (in the case of normally expressed proteins) in panel A, and positive correlation between tissue-specific mutation accessibility (2ε^2^*t*)and tissue protein distance (*D_ti_*) in panel B.

### Relationship between inter-species and inter-duplicate expression divergences

Expression divergence between duplicates has been thought as the major mechanism for duplicate gene preservation (Force et al. 1999; Lynch and Conery 2000; Gu et al. 2005; Kaessmann 2010). For a set of duplicate genes, let *T_dup_*be the mean expression distance between duplicate pairs, or tissue duplicate distance for short. Using 1312 duplicate pairs that were duplicated before the human-mouse split, Gu and Su (2007) estimated *T_dup_* for each tissue, and found a highly significant correlation between tissue expression distance *E_ti_* and *T_dup_*.Similar to the above analysis,we examined the *W_ti_-T_dup_* correlation and the 2ε^2^*t*-*T_dup_* correlation to further explore the underlying tissue factors (Fig.3). While we observed a highly significant *W_ti_-T_dup_* negative correlation (*r*=‐ 0.95, p<0.001), no correlation between 2ε^2^t and *T_dup_* (*p*> 0.10) was found. These observations suggest that when a tissue allows more inter-species expression divergence, it should also tolerate more extensive expression divergence between duplicated genes. Hence, duplicated genes tend to have more expression divergence in a tissue with relaxed developmental constraint, and *vice versa*.

**Figure 3.**
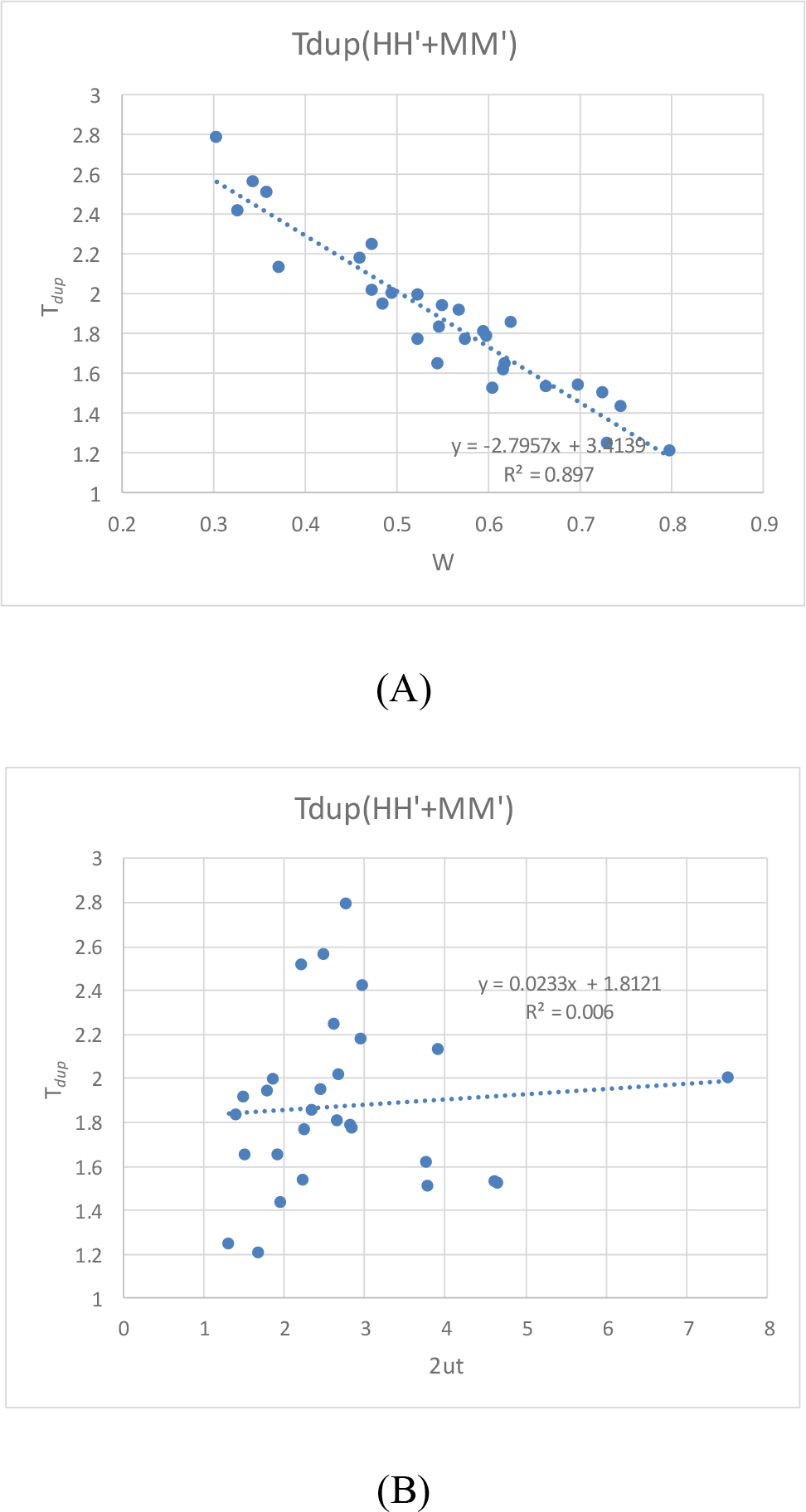
(A) Negative correlation between tissue-specific function constraints (Wti) and tissue duplicate expression distances (*T_dup_*) (*R*^2^=0.87, *p*-value<0.001). (B) No statistically significant correlation between tissue-specific mutation accessibility (2ε^2^*t*)and tissue duplicate expression distances (*T_dup_*) (*p*-value>0.3). Here, *T_dup_* is the average of human and mouse genes.

### Concluding remarks and outlook

In this perspective, we have extended the original tissue-driven hypothesis (Gu and Su 2007) to two sub-hypotheses (tissue-specific functional constraints versus tissue-specific mutational variance), and formulated amodel-based approach to testing these two alternatives. Applying to the human-mouse microarray data, our analysis reveals that both mechanisms may have played nontrivial roles on the tissue-driven evolution, though one should be cautious because of the high noises inherited in the microarray data (Yanai et al. 2004; Yanai et al. 2006). With the rapidincrease of high throughput transciroptome datasets next-generation sequencing (NGS) technologies such as RNA-seq datasets, we speculate that the tissue-driven hypothesis can be further tested in depth as follows.

First, the notion that tissue-specific mutational variance that drives tissue evolution may be shaped by tissue-specific RNA expression profiles (Wang et al 2009; Clark et al. 2011) and epigenetic patterns (She et al. 2009) can be investigated by the genomic analysis combining RNA-seq and other NGS techniques (Morozova et al. 2009). Indeed, previous studies (Tirosh et al. 2006; Cui et al. 2007) have shown that inter-species expression variation may be affected by many gene-specific regulatory factors such as microRNA or TATA box. Second, Arendt (2008) has formulated a conceptual framework witha strong link between cell-type/tissue evolution and expression divergence after gene duplication (Prince and Pickett, 2002; Morkov and Li 2003; Huminiecki and Wolfe 2004; Gu et al. 2005). For instance, high constraint on inter-species expression variation for duplicate genes expressed in the central nerve system (CNS) may be the outcome after arapid evolution toward CNS-specific genes. Third, the relationship between tissue-driven hypothesis and tissue expression broadness can be studied more rigorously because RNA-seq has no the severe cross-hybridization problem that occurred in microarrays.Forth, it would be interesting to test whether the tissue-driven hypothesis still holds for intra-species expression variation such as in the human population (Pickell et al. 2010). And finally, with the availability of RNA-seq datasets in diverse tissue and organisms, we are able to formulate a phylogeny-based framework to study the tissue expression, which would be more powerful than the two species-based analysis. With the help of a newly-developed statistical method (Gu et al. 2013), we have conducted a preliminary analysis by analyzing six tissues (brain, cerebellum, liver, heart, kidney and testis) in mammals (Brawand et al. 2011) and showed qualitatively the same results as shown above (not shown).

## Acknowledgements

We are grateful to Zhan Zhou for constructive comments in the early version of the manuscript, and Jingqi Zhou for technical assistances.

## BOX-1

### Stabilizing selection model

It has been argued that gene expression may be optimized by natural selection (Bedford and Hartl 2009). Gu and Su (2007) invoked the stabilizing selection model (Hansen 1997) to describe the tissue-specific selection constraint on the expression divergence; though many other models were proposed (Gu 2004; Eng et al. 2009). For a gene expressed in a certain tissue (*ti*), the stabilizing selection on the expression level *x* follows a Gaussian fitness function *f_ti_*(*x*) = exp[‐*w_ti_*(*x*‐ θ)^2^], where ff is the optimal expressionlevel, *w_ti_* is the coefficient of stabilizing selection on gene expression in tissue *ti*; a large *w_ti_*means a strong selection pressure, and vice versa. Under this model, the evolution of tissue expression follows an Ornstein-Uhlenback (OU) process, based on which the tissue expression distance is given by 
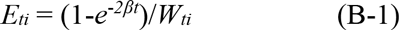

where *W_ti_*=2*N_e_ w_ti_* is the strength of stabilizing selection against the expression divergence, and *N_e_* is the effective population size; β=*W_ti_*ε^2^ is decay-rate of expressi divergence, and ε^2^ is the mutational variance.

### Tissue-specific function-constraint

The hypothesis of tissue-specific constraint postulates that, in multicellular organisms, tissue factors may play important roles in the functional constraint on the rate of expression evolution. As shown in Eq.(1), this effect can be characterized by the tissue-specific parameter *W_ti_*. First, *W_ti_* measures the strength of stabilizing selection on expression evolution. And second, *W_ti_* determines the saturated tissue expression distance, i.e., when *t*→∞, *E_ti_*→ 1/*W_ti_* Hence, the tissue-driven hypothesis predicts that, generally, the parameter *W_ti_* varies among tissues.

### Tissue-specific mutation-accessibility

One of most-exciting findings in genome sciences is that genome-wide epigenetic patterns, such as DNA methylation, histone modification or miRNAs, may have played fundamental roles in shaping the tissue-specific expression profiles in multicellular organisms. It raises the possibility that the among-tissue variation of expression divergence between species could be considerably affected by these epigenetic factors. Tentatively, one may call this phenomenon tissue-specific mutational accessibility, predictinga variation of mutational variance (ε^2^) among tissues.

### Distinguish between two tissue-specific mechanisms

In Eq.(1), we have only one observation (*E_ti_*) but two unknown parameters. To solve this problem, we need an additional equation that can be derived under the assumption of stationary OU process, which means that the expression variance remains invariant during the course of expression evolution. Let *R_ti_* be the coefficient of expression correlation between two species of the same tissue. Under the stationary assumption, it is given by 
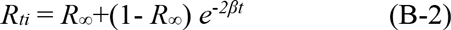

where *R*_∞_ is the coefficient of expression correlation as *t*→∞. In the case of *R*_∞_= 0, one caneasily estimate *W_ti_* from two observations *E_ti_* and *Rti*, that is, by replacing *e*^‐2‐t^ in E q.(B-1) with *R_ti_* according to Eq. (2), we have *W_ti_* = (1‐ *R_ti_*)/*E_ti_*. Meanwhile, from Eq.(B-2) one can estimate 2β*t*=ln *R_ti_*, which leading to the estimation of 2ε^2^*t*=(ln *R_ti_*)/*W_ti_*.

